# RAB5 NUCLEOTIDE BINDING PROMOTES β-OXIDATION TO FUEL HEPATOCELLULAR CARCINOMA CELL PROLIFERATION

**DOI:** 10.1101/2025.08.20.670915

**Authors:** Kelly O. Otakhor, Mohd Ali Abbas Zaidi, Rebecca Oberley-Deegan, Moorthy Ponnusamy, Micah B. Schott

## Abstract

Altered lipid metabolism and lipid droplet (LD) dynamics are hallmark features of hepatocellular carcinoma (HCC) subtypes, but the molecular mechanisms governing LD trafficking and catabolism in HCC cells remain unclear. The small GTPase Rab5, a key regulator of early endosomal dynamics, has been observed to localize to the surface of LDs, suggesting it may play a role in LD turnover. However, the regulation of Rab5-LD interactions and its functional consequences in HCC cell metabolism and proliferation have not been elucidated. In this study, we explored the role of Rab5 in governing LD homeostasis and its impact on HCC cell proliferation. We found that the GTP-bound (Q79L), active form of Rab5 exhibited increased association with LDs compared to the GDP-bound, inactive mutant (S34N). Nutrient starvation enhanced Rab5 GTP-loading and its recruitment to LDs, indicating that Rab5’s GTPase cycle regulates its LD localization. Importantly, inhibition of Rab5 GTP-binding impaired LD catabolism, reduced mitochondrial oxidative phosphorylation, and significantly impaired HCC cell proliferation. Transcriptomic analyses further revealed that RAB5 is significantly overexpressed in HCC patient samples, and this overexpression correlated with poorer overall survival. These findings demonstrate that Rab5’s GTPase cycle is a critical regulator of LD dynamics in HCC cells, governing LD turnover to sustain mitochondrial energy production and support cancer cell proliferation. Targeting the Rab5-mediated regulation of LD metabolism may represent a novel therapeutic strategy to disrupt the metabolic adaptations that fuel liver cancer progression.

## Introduction

Hepatocellular carcinoma (HCC) is currently the fourth leading cause of cancer-related deaths worldwide, characterized by a poor 5-year survival rate largely due to its aggressive clinical course and limited effective treatment options^1–4^. While several targeted therapies have emerged, therapeutic resistance and high recurrence rates continue to challenge long-term disease management^1,3^. In recent years, the rising global prevalence of obesity and excessive alcohol intake has significantly contributed to the surge in HCC incidence^5–8^. As a result, both metabolic dysfunction-associated steatotic liver disease (MASLD) and metabolic dysfunction-associated steatohepatitis (MASH) are rapidly becoming the predominant underlying causes of liver cancer, even in the absence of viral hepatitis^9,10^. Despite the well-established link between steatotic liver disease and HCC development ^11^, the role of lipid droplets (LDs), which are cytosolic organelles that store neutral lipids such as triacylglycerol (TAG) and cholesterol ester (CE), in promoting tumor progression remains underexplored. This study aims to investigate how LD catabolism supports tumor cell proliferation and contributes to the metabolic demands of HCC.

Fatty acid metabolism represents a critical mode of energy homeostasis in a subset of HCC tumors^12^. A central driver of this metabolic adaptation is lipophagy, a selective form of autophagy in which LDs are trafficked to lysosomes for degradation. This process, conserved across species from yeast to mammals, has been documented in many cell types including hepatocytes and macrophages^13–18^. Within the acidic lysosome, lysosomal acid lipase (LAL) hydrolyzes neutral lipids, liberating free fatty acids that can be directed toward mitochondrial β-oxidation or utilized as building blocks for membrane biosynthesis^11,19^. While the enzymatic steps of lipid degradation are well characterized, the upstream processes that govern LD recognition, intracellular trafficking, and lysosomal targeting remain poorly defined. Emerging evidence indicates that cancer cells exploit lipophagy to sustain energy homeostasis and support unchecked proliferation^19,20^. However, in the context of HCC, the molecular regulators orchestrating LD mobilization through this pathway are still largely unknown, highlighting a critical gap in our understanding of tumor metabolism.

Multiple proteomic analyses of LDs have identified a variety of Rab GTPases enriched on the LD surface^21–25^. While Rab proteins are broadly recognized as key regulators of intracellular vesicle trafficking^26^, the specific roles of many Rab family members in LD dynamics remain largely unexplored. Among them, Rab5 stands out for its well-established function in early endosomal trafficking and membrane transport^27–29^. Its activity is tightly regulated through a cycle of GDP-(inactive) and GTP-bound (active) states, coordinated by guanine nucleotide exchange factors (GEFs), GTPase-activating proteins (GAPs), and GDP dissociation inhibitors^29,30^. Once activated, GTP-bound Rab5 recruits its effector proteins such as early endosome Antigen 1 (EEA1), orchestrating the spatial and temporal control of membrane trafficking events^31^. While traditionally studied in the context of endocytosis^32,33^, emerging evidence suggests that Rab5 has broader roles in organelle communication and autophagy^34–38^. Notably, Rab5 has been observed on the surface of LDs^34^. Our recent work has shown that Rab5 localizes to LDs and participates in LD trafficking to lysosomes, implicating it as an important regulator of lipophagy^39^. However, how Rab5 is recruited to LDs, how its GTP/GDP binding activity modulates this process, and whether it plays a direct role in LD turnover in HCC remain open questions.

In this study, we integrate super-resolution microscopy, biochemical assays, and Seahorse-based metabolic profiling to define a mechanistic model of Rab5-mediated lipophagy in HCC. Our findings reveal that Rab5 not only localizes to LDs but is also functionally activated during nutrient starvation to promote LD degradation. Inhibition of Rab5 impairs lipid catabolism, disrupts cellular energy homeostasis, and attenuates HCC cell proliferation. Together, these results position Rab5 as a critical regulator of lipophagy-driven metabolic adaptation in HCC, with potential therapeutic implications.

## Material and method

### Cell Culture and Reagents

Hep3B cells were cultured in Minimum Essential Medium (MEM) containing Earle’s salts and L-glutamine, supplemented with 1 mM sodium pyruvate, non-essential amino acids, and 0.075% (w/v) sodium bicarbonate (Corning). Huh7 cells were maintained in Dulbecco’s Modified Eagle Medium (DMEM) supplemented with 10% fetal bovine serum (FBS), 1% penicillin-streptomycin, and 1% L-glutamine. All cells were grown in a humidified incubator at 37 °C with 5% CO₂. The primary antibodies obtained from Sigma-Aldrich and used in this study were anti-Actin (A2066) and anti-myc tag (9B11). In addition, LDs were stained using Oil Red O (O0625), also from Sigma-Aldrich. Antibodies against Rab5 (2143S), Perilipin-2 (95109), and EEA1 (2411S) were purchased from Cell Signaling Technology. Oleic acid (OA, O1008) was also from Sigma-Aldrich. Rab5 activity was inhibited using neoandrographolide (Rab5 inhibitor, Cayman Chemical, #11742). The constitutively active GFP-Rab5 (Q79L) and dominant-negative GFP-Rab5 (S34N) constructs were gifted from Dr. Sergio Grinstein (Addgene plasmids #35140 and #35141, respectively). The GFP-Rab5 was a gift from Marci Scidmore (Addgene plasmid # 49888). Additionally, Myc-tagged Rab5 was cloned in our lab into the pcDNA3.1Myc/His expression vector, and myc-Rab5(Q79L) and myc-Rab5(S34N) mutants were generated using site-directed mutagenesis. All constructs were confirmed by Sanger sequencing. Hep3B cells were transfected at 60-80% confluency using Lipofectamine 2000 (Thermo Fisher Scientific) and harvested 24-48 hours later for downstream assays.

### Fluorescence Microscopy

Cells were rinsed twice with PBS and then fixed in 3% formaldehyde for 10 minutes at room temperature. To stain LDs, samples were briefly immersed in 60% isopropanol for 30 seconds, incubated in Oil Red O working solution (5 mg/mL in isopropanol, diluted to 60% in water) for 2.5 minutes, and washed again in 60% isopropanol for 30 seconds. Confocal images were captured on a Nikon NSPARC confocal and fluorescent microscope using a 60× oil-immersion objective (NA=1.42). Lipid droplet metrics including droplet number, size, and total area per cell as well as GFP-Rab5 co-localization were quantified in ImageJ. To quantify Rab5 abundance around LDs, LDs were segmented in ImageJ, and corresponding LD regions of interests (ROIs) were converted to peripheral, donut shaped ROIs 0.5 μm in width using the “Make Band” function. Rab5 fluorescence intensity was measured within the band region, and cytoplasmic Rab5 intensity was drawn using a second, hand-drawn ROI. Rab5-LD contact ratio was then calculated by dividing the band intensity of the Rab5 channel (GFP or myc) by the cytoplasmic intensity of the same channel.

### Immunofluorescence Microscopy

Transfected Hep3B cells were seeded onto glass coverslips and treated overnight with 150 μM oleic acid to promote LD accumulation. Following treatment, cells were fixed with 3% formaldehyde at room temperature and subsequently permeabilized using 0.2% saponin. LDs were visualized by staining with Oil Red O (ORO) and myc-Rab5 by immunostaining with the appropriate primary antibodies. After thorough washing, coverslips were mounted using ProLong™ Gold Antifade Reagent containing DAPI (Invitrogen) to label nuclei. Samples were imaged using a Nikon fluorescence microscope equipped with a 60x oil-immersion objective.

### GTP Pull-down Assay

Hep3B cells were harvested and lysed in GTP-Agarose lysis/wash buffer (50 mM Tris-HCl, pH 7.5; 250 mM NaCl; 5 mM Mg acetate; 0.5% Triton X-100; protease inhibitors)^40^. Lysates were clarified by centrifugation (13,000 × g, 10 min, 4 °C). A 50 µL aliquot of each supernatant was reserved as input control. The remaining lysate (∼0.5-1 mg protein) was incubated with 300 µL GTP-agarose beads (Sigma G9768) overnight at 4°C with gentle rotation. Beads were washed in lysis buffer, eluted into Laemmli buffer, and boiled. Samples were resolved by SDS–PAGE and analyzed by western blot.

### Lipid Droplet (LD) Isolation

LDs were isolated from Hep3B cells cultured to approximately 90% confluency in 15 cm dishes (10 dishes per condition). To induce lipid accumulation, cells were treated with 150 µM oleic acid for 16 hours. LD isolation was performed using a modified protocol based on previously published methods^41,42^. Briefly, cells were resuspended in a hypotonic lysis buffer and mechanically lysed using a Dounce homogenizer. The resulting postnuclear supernatant was carefully layered beneath a 30% to 0% OptiPrep density gradient (Sigma, D1556). Following centrifugation at 17,200 rpm for 30 minutes with SW55Ti rotor at 4 °C, the lipid-rich fraction that floated to the top was collected. Isolated lipid droplets were washed and prepared for downstream analysis by western blot. Perilipin-2 (PLIN2) served as a lipid droplet-specific loading control.

### Seahorse Metabolic Analysis

To assess mitochondrial function, adherent cells were seeded at a density of 3 × 10 cells per well in a Seahorse XF96 Cell Culture Microplate (Agilent Technologies), using cell line specific standard growth media. Each experimental group consisted of 10 technical replicates. To ensure even cell distribution, the plates were left undisturbed at room temperature for 20 minutes before being incubated overnight at 37 °C to allow for proper cell attachment. On the following day, the culture medium was replaced with Seahorse XF Base Medium (Phenol Red-free DMEM), and the Cell Mito Stress Test (Agilent) was performed in accordance with the manufacturer’s protocol. After sensor calibration, the assay involved a series of injections with the following compounds to evaluate mitochondrial respiration: oligomycin A (1.5 μM) to inhibit ATP synthase, FCCP (carbonyl cyanide-p-trifluoromethoxyphenylhydrazone; 1.0 μM) to uncouple oxidative phosphorylation, and a mixture of rotenone and antimycin A (0.5 μM) to inhibit complex I and III, respectively. Following the Seahorse assay, cells were lysed using RIPA buffer, and protein concentration was determined via BCA assay. The measured protein content was used to normalize the oxygen consumption rate (OCR) and other metabolic parameters from each well to ensure accurate data comparison.

### Incucyte Live-Cell Assay

Hep3B cells were seeded at a density of 3,000 cells per well in 96-well plates (100 µL/well) using standard growth medium and incubated overnight at 37 °C in a humidified incubator with 5% CO₂. On the following day, the culture medium was aspirated and replaced with 100 µL of treatment medium containing IncuCyte Cytotox Green Reagent for real-time cell health analysis (#4633, Sartorius), prepared according to the manufacturer’s protocol. Cells were treated under the following conditions: DMSO (vehicle control) or NAP. Plates were loaded into the IncuCyte® S3 Live-Cell Analysis System (Essen BioScience) and allowed to equilibrate at 37 °C for 30 minutes before imaging. Time-lapse images were acquired every 3-4 hours using a 10× objective in both phase contrast and green fluorescence channels. Cell proliferation was monitored using phase-contrast imaging to measure confluence over time. All image acquisition and analysis were performed using the IncuCyte integrated cell-by-cell analysis module, allowing for high-content quantification of both viability and growth dynamics at single-cell resolution.

### Colony and Spheroid Formation Assays

To assess both 2D and 3D growth capacities, Hep3B and Huh7 cells were subjected to colony and spheroid formation assays under identical treatment conditions. For the colony formation assay, cells were seeded at 80 cells per well in 24-well plates and allowed to attach for 16 hours. Treatments with 50 µM NAP or DMSO (vehicle control) were applied and maintained for 12 days, with media refreshed as necessary. Colonies were fixed in methanol, stained with 0.5% crystal violet for 5 minutes, rinsed with water, and air-dried. Colony number and size were quantified using ImageJ software software^43^. For the spheroid formation assay, 600 cells per well were seeded into Spherical 5D microwell plates and allowed to aggregate overnight. The resulting cell aggregates were then transferred to CelVivo clinoreactor plates and cultured under identical treatment conditions for 5 days. Spheroids were subsequently collected into standard 6-well plates and imaged for morphological analysis.

## Result

### Rab5 Is Recruited to Lipid Droplets in a GTPase-Dependent Manner in HCC Cells

How Rab5 GTP/GDP binding activity mediates LD recruitment has not been fully elucidated, particularly in HCC. We hypothesized that Rab5’s nucleotide-binding state would dictate its recruitment to LD membranes in HCC cells. To directly assess this, Hep3B human hepatoma cells were transfected with either wild-type (WT) GFP-Rab5, a constitutively active GTP-bound mutant (GFP-Rab5 Q79L), or a dominant-negative GDP-bound mutant (GFP-Rab5 S34N) generated through site directed mutagenesis. It has been previously reported that Rab5 Q79L exhibits persistent activation, leading to enlarged and highly fused endosomal vesicles^44,45^. Thus, we predicted that GFP-Rab5 Q79L would exhibit sustained association with LDs due to enhanced membrane trafficking and endosome-LD interactions. In contrast, the dominant-negative mutant was expected to show diffuse cytoplasmic distribution with minimal LD localization. Wild-type Rab5 was expected to exhibit dynamic localization, with moderate LD localization. We then analyzed the localization of these Rab5 variants relative to LDs stained with ORO using super resolution confocal microscopy (Figure 1A-C). As shown in Figure 1A, cells expressing GFP-Rab5 WT display moderate localization of Rab5 puncta with ORO-labeled LDs (yellow arrows). Notably, expression of “active” GFP-Rab5 Q79L (Figure 1B), resulted in markedly enhanced localization of Rab5 to LDs. This was evident as robust GFP-Rab5 puncta outlining and fully engulfing the LDs. Interestingly, GFP-Rab5 Q79L appeared to preferentially associate with smaller LDs, consistent with the idea that smaller LDs are more amenable to lipophagy^46^. Also, the GFP-Rab5(Q79L) vesicles formed enlarged, highly fused endosomal structures engulfing smaller LDs. Cells expressing the GDP-bound inactive GFP-Rab5 S34N mutant (Figure 1C) also exhibited interaction around LDs, although there was much more prominent cytoplasmic localization compared with WT and QL Rab5.

**Figure 1.**
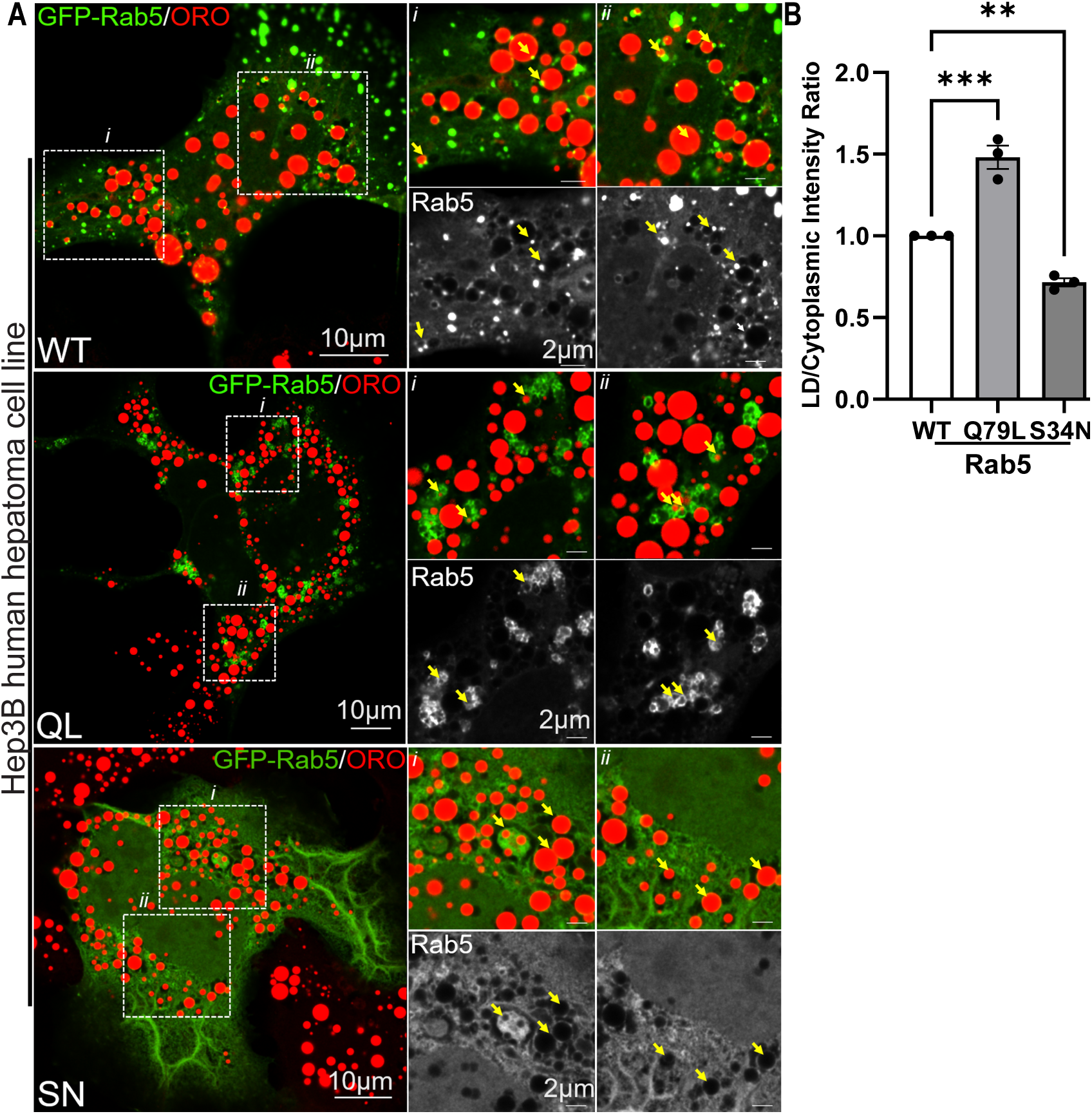
legend: Rab5 Is recruited to lipid droplets (LDs) in a GTPase-dependent manner in Hep3B cells. Super-resolution confocal micrographs of Hep3B human hepatoma cells expressing (A) wild-type Rab5 (GFP-Rab5 WT), (B) constitutively active mutant (GFP-Rab5 Q79L), and (C) dominant-negative mutant (GFP-Rab5 S34N). LDs were stained with Oil Red O (ORO, red) to visualize neutral lipid stores. GFP-Rab5 fluorescence (green) marks Rab5 localization. Each panel contains two high-magnification insets (i and ii): inset i and ii zooms in on Rab5 puncta at LD borders, while below each of this insets shows the grayscale GFP channel alone, highlighting Rab5 signal independent of ORO. In Panel A, yellow arrows indicate discrete punctate Rab5 structures in close proximity to ORO-stained LDs, representing moderate LD association by wild-type Rab5. In Panel B, yellow arrows highlight continuous GFP-Rab5 rings surrounding smaller LDs, showing enhanced recruitment of the GTP-locked Rab5 Q79L to LD surfaces. In Panel C, yellow arrows mark areas where LDs lack adjacent Rab5 signal, and the GFP-Rab5 S34N mutant remains diffusely distributed throughout the cytoplasm, confirming a failure in membrane targeting due to its GDP-bound inactive state. (D) Quantification of Rab5-LD co-localization. The LD/Cytoplasm Intensity Ratio was calculated for each mutant by dividing the average GFP-Rab5 intensity at LD surfaces by the cytoplasmic GFP-Rab5 intensity. Bars represent mean ± SD from 3 independent experiments. Statistical significance was determined using one-way ANOVA with Tukey’s post hoc test. **p<0.01 and ***p<0.001. Scale bars: main images, 10 µm; insets, 2 µm.

Quantification of Rab5-LD: cytoplasmic localization intensity ratios (Figure 1D) showed a ∼1.7-fold increase in Rab5 Q79L-LD association relative to Rab5 WT, while Rab5 S34N displayed a decrease in LD localization compared to Q79L. These data demonstrate that the GTP-bound active form of Rab5 localizes more exclusively to LDs in Hep3B cells, while the GDP-bound Rab5 also localizes but to a lesser degree, suggesting that Rab5-LD interaction is regulated by Rab5’s nucleotide binding state. These findings support a model in which Rab5’s ability to bind GTP enhances its recruitment to LDs. In HCC cells, this places Rab5 activation as a critical regulatory checkpoint for LD catabolism, linking its GTPase cycle directly to the control of lipid homeostasis.

### Enhanced Recruitment of Rab5 to LDs during Lipophagy Stimulation

Next, as nutrient starvation is known to stimulate lipophagy^16,47,48^, we evaluated the effect of Hank’s Balanced Salt Solution (HBSS) starvation on Rab5-LD recruitment as well as Rab5 GTP/GDP binding. Hep3B cells were starved 4hours in HBSS, and GTP-bound Rab5 levels were assessed by pull-down with GTP-binding agarose beads. We observed a >50% increase in Rab5 activity in starved cells compared to cells maintained in complete media (control) (Figure 2A & B), suggesting that Rab5 is active during lipophagy stimulation. Levels of EEA1, a known interactor with active Rab5, trended higher in starved cells, but this was not statistically significant (Figure 2A & B). As an additional control, we observed higher pulldown of active myc-Rab5 Q79L but not the dominant-negative myc-Rab5 S34N using GTP-beads (Figure S1A). These results indicate that Rab5 is not only recruited to LDs during nutrient deprivation but is also activated through GTP loading. This is consistent with our results from Fig. 1A, as well as previous reports showing Rab5 enrichment in LD fractions treated with a non-hydrolysable GTP analog (GTPγS)^34^

**Figure 2.**
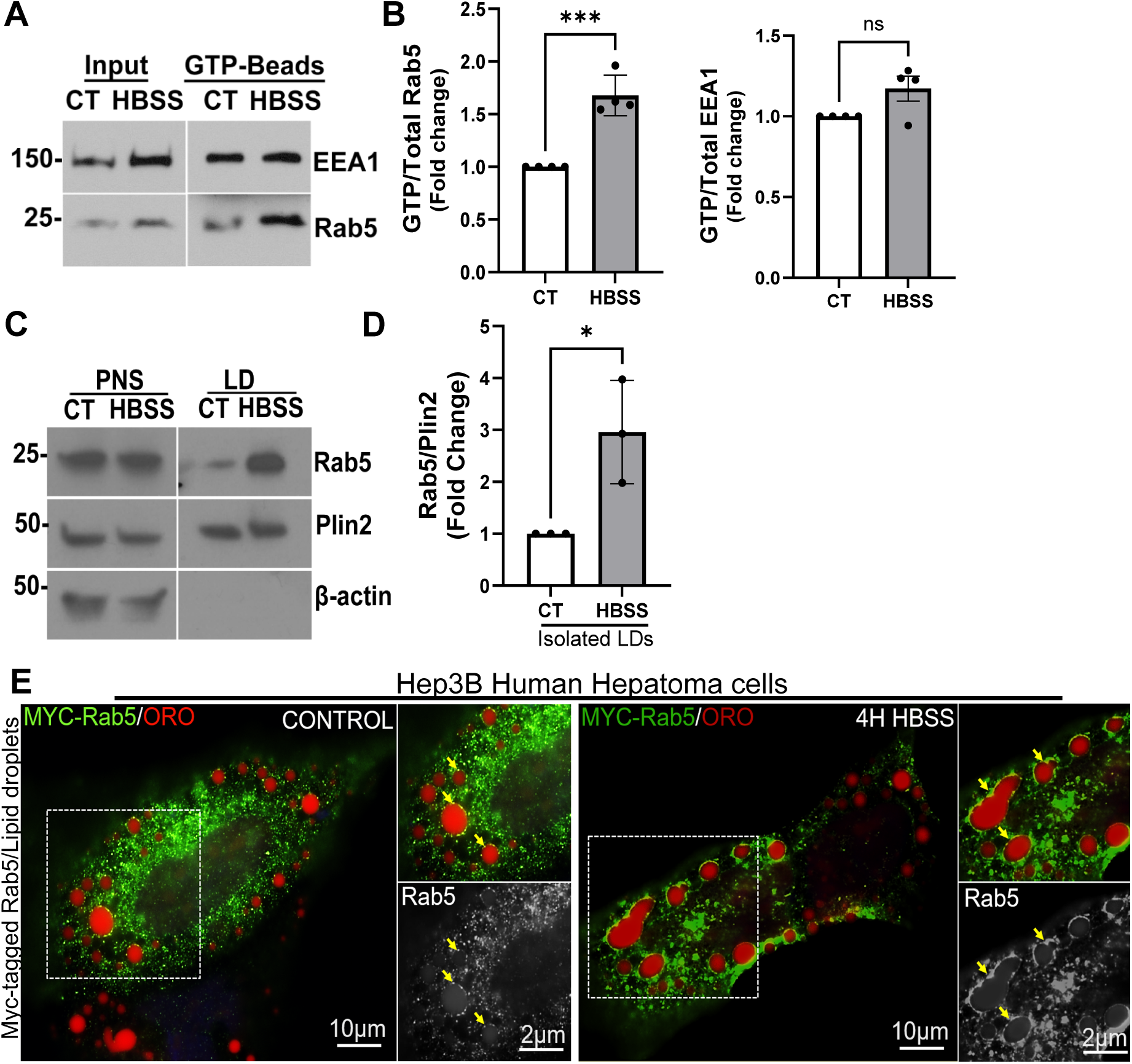
legend: Enhanced recruitment of Rab5 to LDs upon lipophagy stimulation. A) Western blot analysis of Rab5 and EEA1 GTP-pulldown in 4h HBSS and control (complete media) treated condition in Hep3B cells. Note the increase in active Rab5 in the lipophagy stimulated condition (HBSS) compared to control (complete media). B) Quantification of the fold change of GTP-bound/total Rab5 and GTP-bound/total EEA1 ratios from four pull-down experiments (mean ± SD). ***p<0.0001 and p=0.0683 by unpaired two-tailed *t*-test. C) Western blot analysis of LDs isolated from Hep3B cells by density gradient centrifugation show abundant levels of Rab5 in isolated LDs following 4 hours of HBSS starvation compared to the control (complete media). Note: Plin2 served as an LD marker, and absence of β-actin in the LD fraction shows that there was no cytoplasmic contamination. D) Quantification of ratio of Rab5/Plin2 in the biochemically isolated LDs from n=3 independent experiments. p=0.0270 by unpaired two-tailed Student’s t-test; asterisks indicates p<0.05. E) Fluorescence micrograph showing immunofluorescence staining of Hep3B cells expressing Myc-tagged Rab5 (green) maintained in complete media (Control) or subjected to 4 h HBSS starvation, then fixed and stained with Oil Red O (ORO; red) to label LDs. Left panels show a full-cell view under control and starvation conditions (scale bar, 10 μm). Right panel contains one inset zooms in on Rab5 puncta at LD borders, while below the insets shows the grayscale GFP channel alone, highlighting Rab5 signal independent of ORO. (grayscale; scale bar, 2 μm). In control cells, Rab5 puncta are infrequently associated with LDs, whereas HBSS starvation markedly increases the number of Rab5-positive puncta apposed to or encircling LDs (yellow arrowheads), indicating enhanced recruitment of active Rab5 to LD surfaces under nutrient stress.

Given our findings that HBSS increases Rab5 activity (Figure 2A), and that active Rab5 exhibits greater association with LDs (Figure 1B), we sought to determine the effect of HBSS starvation on Rab5 recruitment to LDs. Hep3B cells were first loaded with oleic acid (150μM, 16h) to induce LD formation, then cells were stimulated by acute nutrient deprivation using HBSS for 4 hours. Following this treatment, LDs were isolated by density gradient centrifugation, and Rab5 was assessed by western blot analysis. As shown in Figure 2C-D, Rab5 levels were elevated 2-4 fold in LD fractions in starved cells compared with control cells maintained in complete media, indicating enhanced recruitment of Rab5 to LDs during lipophagy activation. Plin2 served as an LD marker and loading control, while the absence of β-actin in LD fractions confirms the lack of cytoplasmic contaminants. Immunofluorescence microscopy further corroborated the biochemical findings (Figure 2E). In Hep3B cells maintained in complete media, myc-tagged Rab5 (green) exhibited a predominantly diffuse cytoplasmic distribution, with limited localization with ORO-stained LDs (red). In contrast, Rab5 was more predominantly localized around LDs following HBSS-induced starvation as indicated by white arrows. This enhanced localization suggests increased Rab5 recruitment to LDs under nutrient-deprived conditions, consistent with its role in mediating lipophagy^39^. Together, during nutrient starvation, these findings support a key role for Rab5 in lipophagy by initially targeting LDs to coordinate their trafficking and endolysosomal catabolism.

### The Pharmacological Inhibition of Rab5 Activity Decreases LDs Catabolism and GTPase Activity

Building on our findings that Rab5 is both recruited to LDs and activated during nutrient starvation (Figure 1 & 2), we next sought to determine whether Rab5 GTP binding activity is functionally required for LD catabolism. To test this, we utilized the small molecule neoandrographolide (NAP), a diterpenoid derived from *Andrographis paniculata*^49^, which has been previously identified as a selective and potent inhibitor of Rab5 GTPase activity^50^. We first validated the effect of NAP on Rab5 GTP binding state using GTP-agarose beads in Hep3B cells. As expected, treatment with NAP (100μM, 48h) resulted in a >90% reduction Rab5 activity (Figure S2A-B). We next assessed the effect of NAP treatment on LD accumulation in ORO-stained Hep3B cells. As shown in Figure 3A, NAP treatment resulted in a marked increase in both LD number and total area per cell, compared with DMSO-treated controls. Quantitative analysis (Figure 3B) demonstrated a ∼2-3-fold increase in total LD area per cell upon Rab5 inhibition, with no significant change in average LD size, but a ∼3-fold increase in LD number per cell. The effect of NAP was especially visible in oleic acid-treated Hep3B cells, as shown in Figure 3C (note the dramatic accumulation of LDs in NAP vs DMSO-treated cells). Quantitative analysis (Figure 3D) revealed a ∼5-fold increase in LD area per cell, a ∼1.5-fold increase in LD size, and a ∼3-fold increase in LD number per cell following Rab5 inhibition. These results collectively establish that Rab5 activity is required for LD catabolism, and that its pharmacological inhibition leads to impaired LD turnover due to a loss of GTP-bound Rab5.

**Figure 3.**
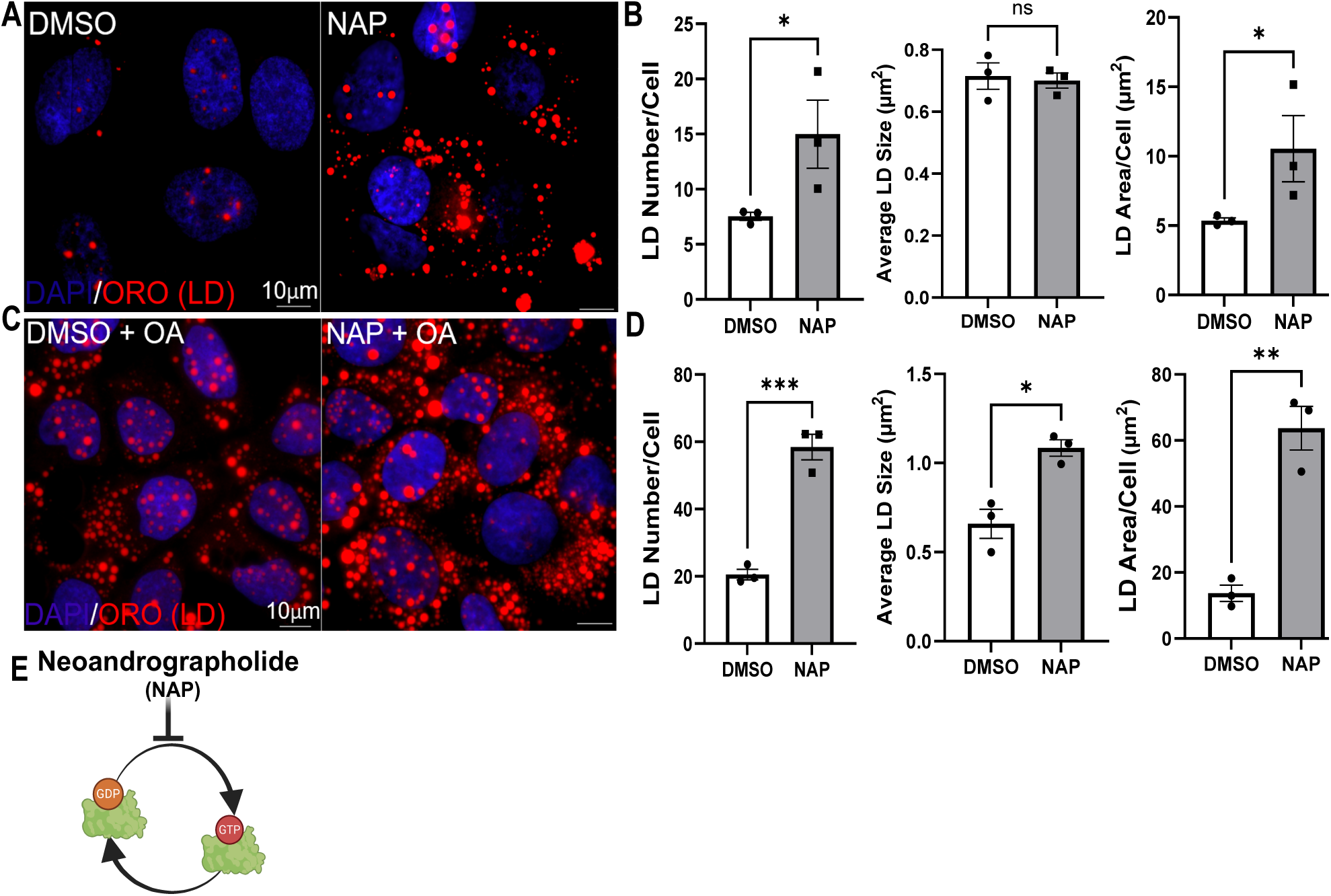
legend Inhibition of Rab5 GTPase activity reduces LD catabolism via its GTPase activity. A) Fluorescent micrograph showing ORO-stained LDs (red) and Dapi stained nucleus from Hep3B cells treated with DMSO (control) and NAP (100µM). B) Bar graphs depict quantification of total LD area/cell, LD size, and LD number/cell (mean ± SD; n=3 experiments; *p<0.05, p=0.3898 unpaired two-tailed Student’s *t*-test). C) Fluorescent micrograph showing ORO-stained LDs (red) and Dapi stained nucleus from Hep3B cells treated with DMSO (control) and NAP (100µM) with oleic acid (OA, 150 µM). D) Bar graphs depict quantification of total LD area/cell, LD size, and LD number/cell. (mean ± SD; n=3 experiments; *p<0.05, **p<0.001, ***p<0.0001, unpaired two-tailed Student’s *t*-test). E) Cartoon depicting NAP alteration of Rab5 GTP binding.

### Inhibition of Rab5 GTP/GDP Binding Activity Impairs Mitochondrial Respiration in HCC Cell

During nutrient deprivation, cancer cells can switch to utilization of fatty acids (FAs) as fuel for mitochondrial oxidative phosphorylation to maintain ATP production. Given the essential role of Rab5 in coordinating LD trafficking and catabolism under nutrient starvation, we reasoned that Rab5-GDP/GTP binding activity could directly impact mitochondrial energy production under this condition. To assess this relationship, we evaluated mitochondrial function in Rab5-inhibited Hep3B cells using the Seahorse Mito Stress Test, which provides real-time measurement of oxidative phosphorylation and allows different stressors to be added during the assay to challenge the mitochondria such as oligomycin (inhibits ATP synthase), FCCP (uncouples the electron transport chain), and rotenone/antimycin A (blocks complex I/III) allowing us to derive multiple key parameters such as basal respiration, ATP-linked respiration, maximal respiration, non-mitochondria respiration and spare respiratory capacity. Of particular interest to us was the spare (or reserve) respiratory capacity, a key metric that reflects the ability of mitochondria to respond to increased energy demands by ramping up ATP production. This parameter represents the extra oxidative capacity available beyond basal respiration and is considered a marker of mitochondrial fitness and metabolic flexibility^51,52^. Previous studies have demonstrated that cells with high spare respiratory capacity can more efficiently generate ATP under stress conditions ^51^.

Consistent with this, we found that Rab5 inhibition significantly suppressed oxygen consumption rate (OCR) in both complete media (CM) and nutrient-deprived (HBSS) conditions. In complete media (Figure 4A-B), NAP-treated cells showed a pronounced reduction in both basal respiration and spare respiratory capacity, indicating a loss of mitochondrial responsiveness and energy-generating potential. Under starvation conditions (4-hour HBSS pretreatment, Figure 4C-D), Rab5 inhibition continued to suppress OCR, further reducing both basal and spare capacity, reinforcing the idea that Rab5 is critical for sustaining mitochondrial oxidative metabolism, especially during lipophagy stimulation. Taken together, these data demonstrate that Rab5 inhibition impairs energy production, leading to metabolic stress in HCC cells.

**Figure 4.**
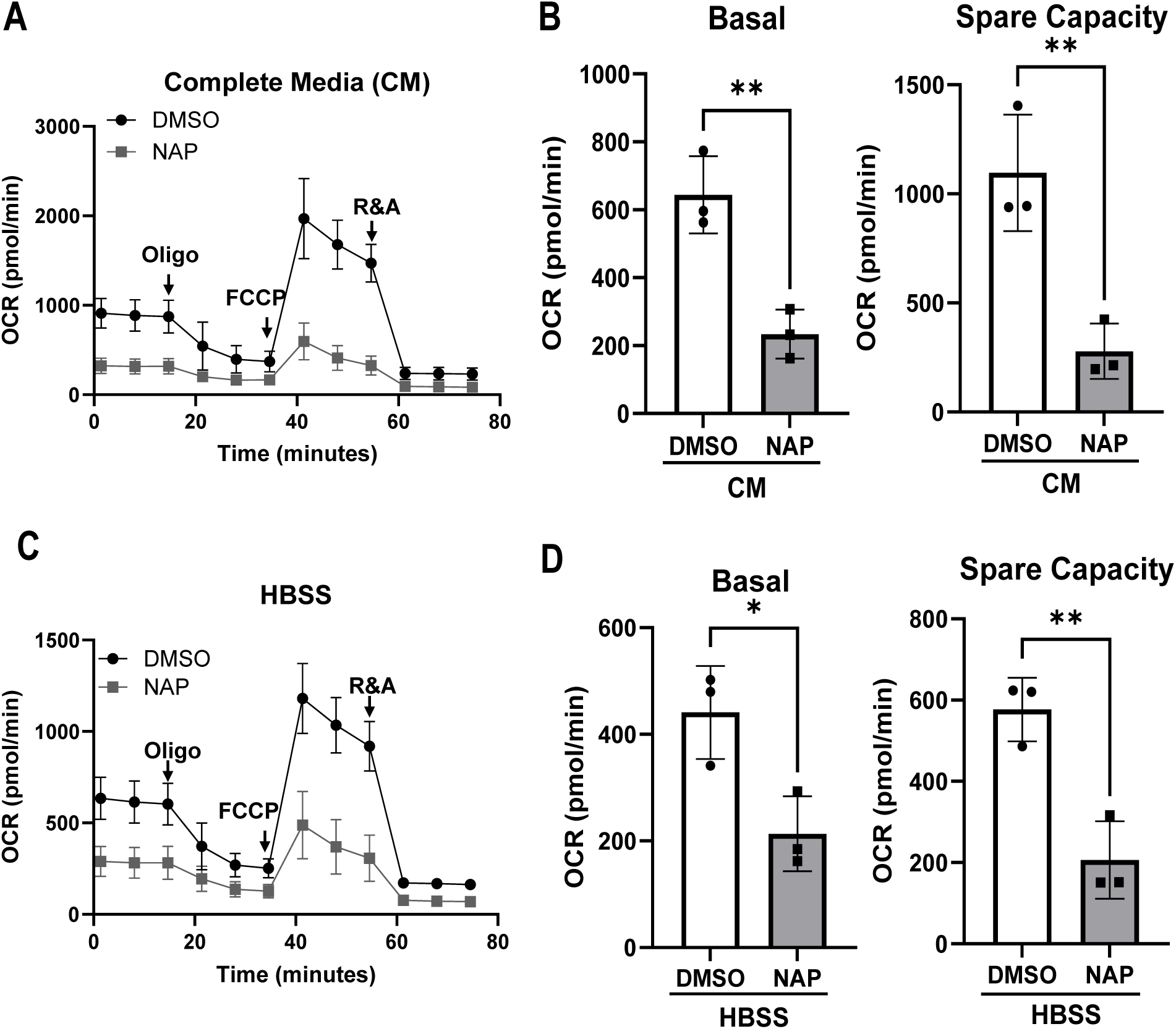
legend: Inhibition of Rab5 decreases HCC energy homeostasis. A) Seahorse metabolic analysis showing NAP (100 μM, 48 h) inhibition of Rab5 decreases oxygen consumption rate (OCR) compared to control in complete media. n=3 independent experiments. B) Quantification of basal and spare respiratory capacity in complete media following treatment with NAP. n=3 independent experiments, (mean ± SD). **p<0.01 by unpaired two-tailed *t*-test. C) Seahorse metabolic analysis showing NAP (100 μM, 48 h), inhibitor of Rab5, during 4 hours HBSS starvation decreases oxygen consumption rate (OCR) compared to control (DMSO) n=3 independent experiments. D) Quantification of basal and spare respiratory capacity in 4hours HBSS media following treatment with NAP. n=3 independent experiments, (mean ± SD). *p<0.05, **p<0.01 by unpaired two-tailed *t*-test.

### Inhibition of Rab5 GTP/GDP Binding Activity Decreases HCC Cell Proliferation

To investigate the functional consequences of Rab5 GTP/GDP binding activity inhibition on tumorigenic potential, we assessed the ability of HCC cells to form colonies and spheroids upon disruption of Rab5 activity (Figure 5A-D). HCC cell lines (Hep3B and Huh7) were subjected to colony formation and 3D spheroid assays following treatment with NAP. In the colony formation assay (Figure 5A), untreated control cells exhibited robust colony growth, forming an average of ∼40-55 colonies per well in both Hep3B and Huh7 lines. In contrast, treatment with NAP significantly impaired colony formation. Quantification (Figure 5B) revealed that NAP reduced colony numbers by ∼80-90% compared to untreated controls, highlighting the essential role Rab5 activity in sustaining clonal proliferation of HCC cells.

**Figure 5.**
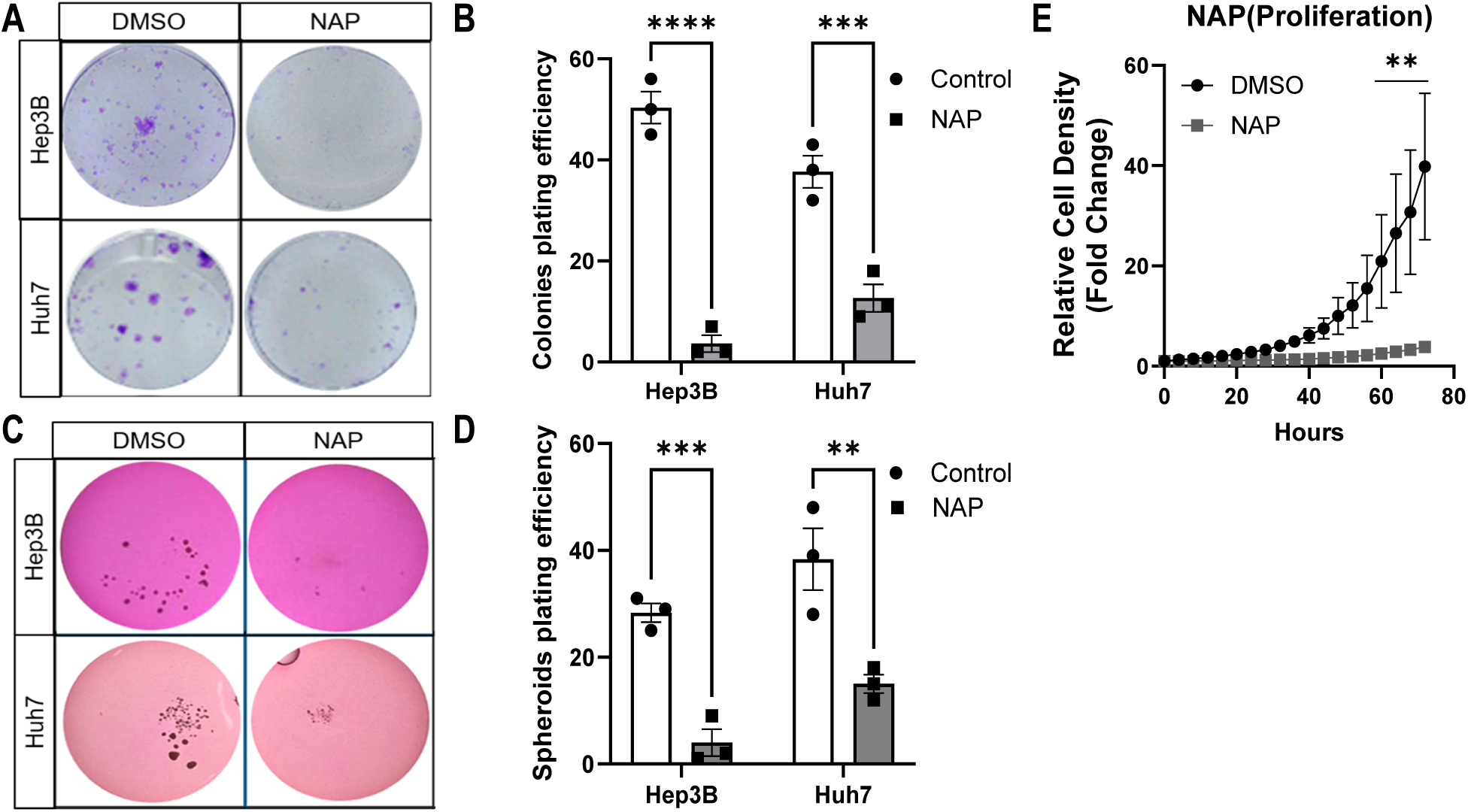
legend: NAP treatment decreases HCC cell proliferation HCC. A) Representative images showing colony formation assay of Huh7 and Hep3B cell treated with NAP (50μM) for 12 days. B) Quantification of colony formation assay of Huh7 and Hep3B cell treated with NAP (50μM) for 12 days. n=3 independent experiments. Statistical significance was determined using two-way ANOVA. ***p<0.0004 and ****p<0.0001. C) Spheroid formation assay of Huh7 and Hep3B cells treated with NAP (50μM) for 5 days. n=3 independent experiments. D) Quantification of spheroid formation assay of Huh7 and Hep3B cells treated with NAP (50μM) for 5 days. n=3 independent experiments. Statistical significance was determined using two-way ANOVA. **p<0.005 and ***p<0.001. E) Quantification of IncuCyte live-cell imaging of Hep3B cells treated with DMSO (vehicle control) or NAP (100μM), and relative cell density (fold change) was monitored over 72 hours. Data represents mean ± SD of n=3 biological replicates.

To evaluate the impact on 3D growth, we performed a spheroid formation assay (Figure 7C). Consistent with the 2D colony results, NAP markedly reduced spheroid size and number in Hep3B and Huh7 cells as compared to controls. While control cells formed large, dense spheroids indicative of high proliferative and metabolic capacity, lipophagy-inhibited cells generated either poorly formed or fewer spheroids, reflecting impaired self-renewal and growth potential. Together, these findings suggest that Rab5-driven lipophagy may support not only energy homeostasis (Figures 5 and 6), but also the clonogenic and tumorigenic potential of HCC cells. The consistent suppression of both colony and spheroid formation upon Rab5 inhibition supports the model wherein Rab5-mediated LD catabolism fuels key oncogenic programs, including metabolic adaptation and sustained proliferation.

**Figure 6.**
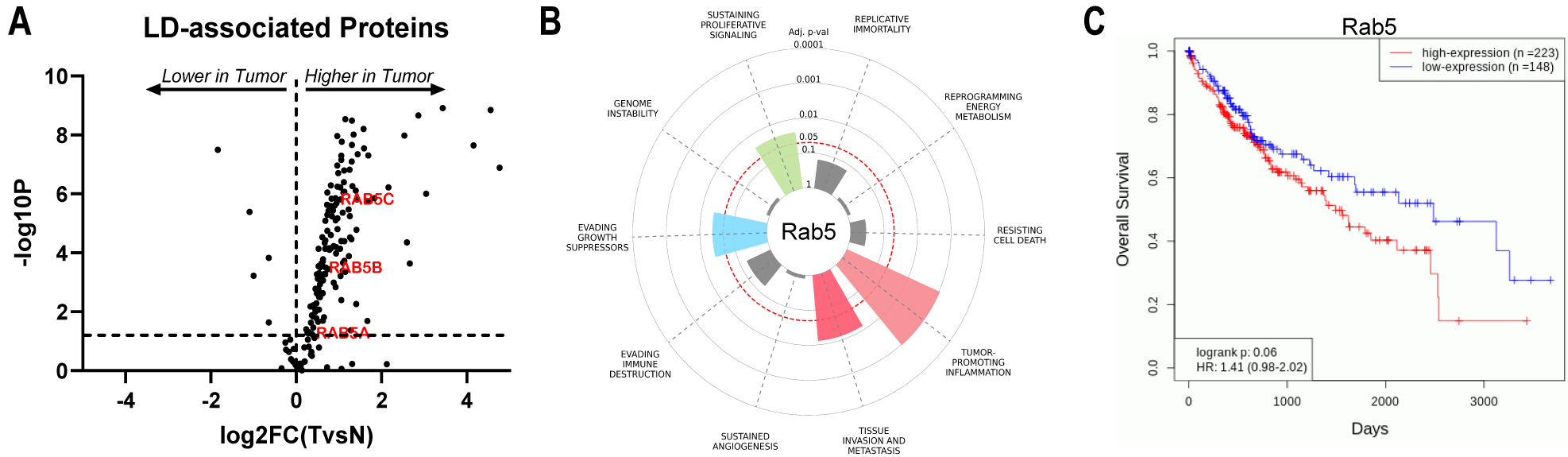
legend: Rab5 is upregulated in HCC and associated with cancer hallmarks. (A) Volcano plot from TNMplot showing expression of known LD-associated proteins including Rab5A/B/C in HCC tumors vs. adjacent normal tissues from the same patient. Log₂ fold-change is on the x-axis and –log₁₀ p on the y-axis. Dashed lines at ±1 (2-fold change) and –log₁₀ p = 1.30 (p=0.05) define significance cutoffs; Rab5A, Rab5B, and Rab5C are notably upregulated in tumors. (B) Cancer hallmark gene set enrichment analysis based on Menyhart et al.^54^, showing Rab5-associated genes enriched in proliferation, invasion, and evading growth suppressors. (C) Kaplan-Meier survival analysis from TCGA-LIHC dataset stratified by Rab5 expression. High Rab5 expression correlates with poorer overall survival (log-rank test, p=0.06).

**Figure 7.**
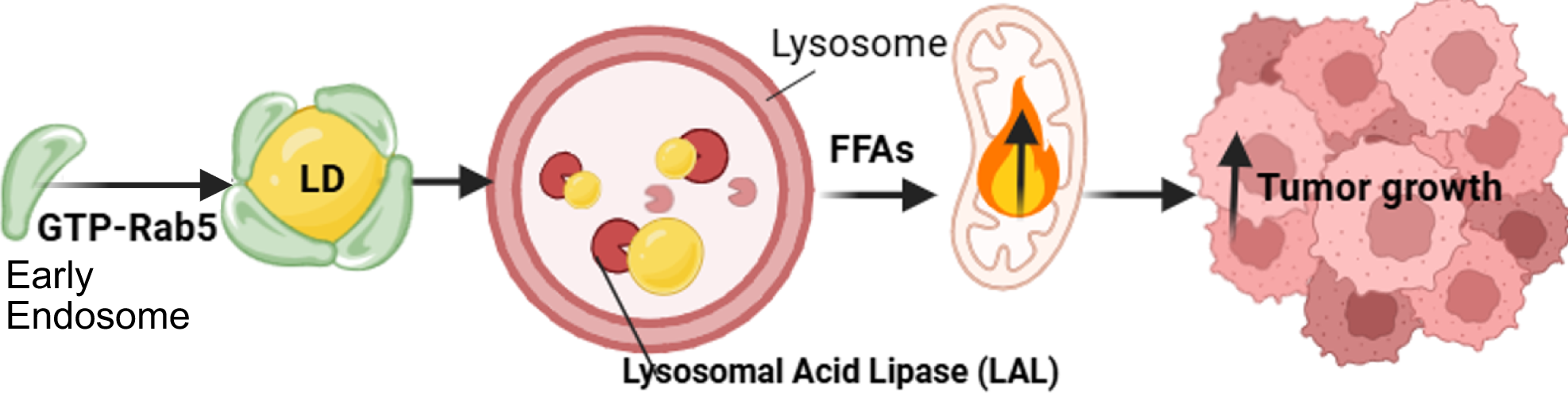
legend: Working model depicting Rab5-mediated LD catabolism in HCC. Under nutrient deprivation, Rab5 is activated by exchanging its bound GDP for GTP and translocates to the LD surface, where GTP-Rab5 drives tethering and fusion with lysosomes. Inside the lysosome, lysosomal acid lipase (LAL) degrades LD, releasing free fatty acids (FFAs). These FFAs are then shuttled into mitochondria for β-oxidation, generating the ATP that fuels HCC cell proliferation and survival.

To further assess the impact of Rab5 inhibition on HCC cell growth, we performed a real-time proliferation assay using IncuCyte live-cell imaging for 72 hours (Figure 5E). Cells treated with NAP showed near-complete suppression of proliferation compared to DMSO controls, with minimal increase in cell density over time. These findings indicate that Rab5 activity is not only essential for maintaining lipid metabolism and mitochondrial function, but also for driving HCC cell expansion.

### Rab5 Is Overexpression in HCC Indicates Adverse Clinical Outcomes

To investigate the differential expression of Rab5 isoforms (A, B and C) and LD-associated proteins in HCC, we conducted a transcriptomic analysis using publicly available datasets from GEO, GTEx, TCGA, and TARGET. These datasets collectively include gene expression profiles from both tumors and adjacent normal liver tissues, providing a comprehensive view of transcriptional changes associated with liver tumorigenesis. A curated gene list of known LD-associated proteins was compiled to assess their expression dynamics in HCC. Analysis of this dataset revealed that RAB5A, RAB5B, and RAB5C were significantly upregulated in HCC tumors compared to adjacent normal tissue (Figure 6A, Table S1)^25,53^.

To determine which core oncogenic processes are most closely associated with elevated Rab5 expression in cancer, we utilized a previously curated list of cancer hallmark gene sets derived from the integrated cancer hallmark gene set assembled from catalogue of somatic mutations in cancer (COSMIC) and curated literature sources for enrichment analysis using the ten canonical “Hallmarks of Cancer” (Table S1)^54^. For each hallmark, we calculated how many Rab5 genes overlapped with the hallmark gene set, the statistical significance of that overlap (P-value and FDR-corrected Adjusted P-value), and the fold-enrichment (Odds Ratio)^54^. The circular bar chart (Figure 6B) and hallmark overrepresentation table (Table S2) highlights strong enrichment of Rab5-associated genes in tumor-promoting inflammation, invasion and metastasis, evasion of growth suppressors, and sustained proliferative signaling. Conversely, little to no enrichment is seen for genome Instability, sustained angiogenesis, and reprogramming energy metabolism, suggesting Rab5’s principal roles in proliferative and invasive pathways in cancer.

We next assessed the prognostic significance of Rab5 overexpression, we next performed Kaplan-Meier survival analysis, which incorporates clinical outcome data from over 10,000 cancer patients in the TCGA study and corresponding normal tissue data from the GTEx project. The results (Figure 6C) showed that patients with high Rab5 expression exhibited poorer overall survival compared to those with low expression in liver hepatocellular carcinoma (LIHC), indicating that Rab5 overexpression is a negative prognostic biomarker in HCC and underscoring the prognostic potential of Rab5 isoforms in HCC^55,56^.

Overall, these integrated analyses encompassing differential expression, hallmark enrichment, and survival correlations demonstrate that Rab5 are consistently upregulated in HCC, are implicated in core oncogenic processes (particularly proliferation and invasion), and portend worse patient outcomes. This supports Rab5 as both a mechanistic driver of aggressive liver cancer phenotypes and a potential target for therapeutic intervention.

## Discussion

LD catabolism provides essential fuel for tumor growth and survival under metabolic stress across multiple cancer types, yet the molecular regulators and trafficking pathways that govern LD degradation in HCC remain poorly characterized. The data presented here defines a mechanism by which nutrient deprivation directly activates Rab5’s nucleotide binding, driving its recruitment to LDs and initiation of their degradation, potentially via microlipophagy. This process liberates free fatty acids to sustain mitochondrial β-oxidation and support HCC proliferation (Figure 7). We further show that pharmacological blockade of Rab5’s GTP-bound state with neoandrographolide (NAP) perturbs LD clearance, diminishes mitochondrial respiration, and abrogates both 2D colony and 3D spheroid growth. We propose that Rab5 is activated upon low nutrient availability and responds by directing LDs to lysosomes, potentially via microlipophagy, to fuel HCC growth.

Nutrient deprivation activates several Rab GTPases as frontline responders to metabolic stress. Our group previously demonstrated that LD trafficking to lysosomes relies on Rab5-dependent endosomal pathways rather than the macroautophagy machinery (Atg5/FIP200)^39^. Mechanistically, we showed that the GTP-bound Rab5 localize to the LD, whereas GDP-bound Rab5 is more cytosolic signifying less or no localization (Figure 1). This mirrors classical Rab5 biochemistry in which *in vitro* reconstitution experiments showed that Rab GDP dissociation inhibitor (RabGDI) extracts inactive Rab5 from LDs and only the GTP-loaded form can re-associate and recruit its effectors like EEA1 to enhance membrane attachment^34,57^. Under HBSS-induced starvation, we observe a dramatic increase in Rab5 GTP loading and its recruitment to LDs (Figure 2), echoing reports of Rab5 activation during serum withdrawal promotes PI3K-Beclin1 complex formation and autophagy initiation^58,59^. Amino acid withdrawal increases the colocalization of BODIPY-stained LDs with Rab5-positive endosomal compartments, indicating that nutrient stress brings Rab5-bearing membranes into close apposition with LDs^60^. Interestingly, other Rab GTPases are also activated during nutrient starvation to stimulate lipophagy. For example, Rab7 is strongly GTP-loaded under starvation to orchestrate lysosome-mediated LD degradation^61^ and Rab10 is similarly activated to facilitate LD engulfment during lipophagy^48^. Reinforcing the evolutionary conservation of nutrient starvation to activate Rab5, yeast cells transcriptionally upregulate the Rab5 paralog Ypt53 under nitrogen deprivation to boost endosomal trafficking and vacuolar activity^62^.

In cancer, where the tumor microenvironment is marked by limited nutrients and oxygen, cancer cells hijack the cellular starvation response to activate LD catabolism and sustain their metabolic demands^63–65^. Mounting evidence indicates that lipophagy is a major route for LD degradation in hepatocytes^46,66–68^ under nutrient starvation or stress^66,68^. In support of this, liver-specific Rab5 knockdown in mice leads to marked LD accumulation and impaired mitochondrial β-oxidation^69^, underscoring Rab5’s essential role in efficient lipid catabolism and mobilization. Consistent with these observations, we showed that disrupting Rab5’s nucleotide cycle using NAP halts LD clearance (Figure 3), demonstrating that GTP-bound Rab5 and recruitment of its effector EEA1 (Supporting Information 2A-B) is not merely correlative but functionally essential for LD trafficking and catabolism. Despite Rab5 inhibition, total EEA1 levels remained largely unchanged (Figure S2A-B), indicating that NAP disrupts EEA1’s localization rather than its expression or stability. One explanation is that, without GTP-loaded Rab5, EEA1 simply remains in the cytosol or associates with non-LD endosomes instead of docking at LDs. Alternatively, a minimal pool of Rab5-GTP may suffice to maintain basal EEA1 expression but not its efficient recruitment to LD membranes. Additionally, we reveal that Rab5-mediated LD degradation is indispensable for fatty acid driven ATP production in Hep3B cell (Figure 4A-D). Under both nutrient-replete and nutrient-deprived conditions, Rab5 inhibition with NAP causes a dramatic collapse of mitochondrial β-oxidation demonstrating that, even when external fuels are abundant, HCC cells depend on Rab5-driven LD catabolism to maximize mitochondrial oxidation of intracellular lipid stores. By blocking Rab5, we unmask this critical reliance on LD catabolism for meeting energetic demands in Hep3B cell. Functionally, pharmacologic blockade of Rab5 with NAP profoundly impairs cell proliferation (Figure 5A-E). NAP has been shown to suppress growth of lung cancer cells through competing with Rab5 GTP/GDP binding sites and preventing the GTPase from cycling into its active state^50^, underscoring its potential as a broad-spectrum anticancer agent. Given these impacts on HCC cell metabolism and proliferation, NAP emerges as a promising therapeutic candidate for targeting Rab5 in HCC.

Cancer cells hijack a broader remodeling of the Rab GTPase network to meet heightened demands for membrane trafficking, nutrient scavenging, and survival. Our transcriptomic analyses (Figure 6A-C) place Rab5 among the upregulated genes in HCC tumors, with high Rab5 expression correlating with poor patient survival. It is possible that upregulation of Rab5 enhances cell proliferation^70–72^, promotes integrin recycling and resistance to anoikis^73^, and accelerates focal-adhesion turnover to facilitate migration and invasion^74^. While the exact triggers of Rab5 overexpression in HCC remain undefined, it is potentially driven by a convergence of oncogenic growth-factor signaling, hypoxia-induced pathways, and nutrient-stress sensors that together enhance Rab5 transcription and GTP-loading in tumor cells.

One of the key limitations of our study is that we cannot yet distinguish whether Rab5 binds directly to the LD monolayer or instead localizes indirectly via early endosomal membranes that dock onto LDs. Electron microscopy work has shown that transferrinl1HRP labeled early endosomal tubules form intimate contacts with LD surfaces ^35^, suggesting that Rab5 may simply reside on these endosomal membranes rather than embedding in the LD monolayer itself. Resolving this will require higherl1resolution imaging or biochemical fractionation approaches that can clearly separate endosomal from LD membranes. Also, the specific upstream nutrient-sensing kinases or GEFs/GAPs responsible for modulating Rab5 remain unknown, but emerging evidence points to energy-sensing kinases as likely intermediaries. For instance, in muscle cells, AMPK phosphorylates a Rab5-specific GAP to increase Rab5 activity and promote GLUT4 translocation^75^, raising the possibility that similar stress-activated kinases may govern Rab5 activation in hepatocytes.

Despite clear evidence that both HBSS starvation and NAP treatment inhibit mitochondrial β-oxidation and potentially block LD clearance in HCC cells, several caveats temper our interpretation. First, although NAP is reported to inhibit Rab5 GTPase activity, it may also disrupt other Rab5-dependent processes or have off target effects such as clathrin-mediated endocytosis, macropinocytic nutrient uptake, general autophagy initiation, multivesicular body biogenesis, and integrin recycling thereby broadly impairing cellular signaling, cell migration and adhesion^37,59,73,75–82^. Second, acute HBSS treatment deprives cells of multiple nutrients and can nonspecifically suppress mitochondrial respiration and activate global autophagy programs, making it challenging to isolate Rab5-specific lipophagic effects. Third, while lipophagy is likely the dominant LD catabolic route in hepatocytes, we have not directly measured autophagic flux on LDs (for example, using LD-targeted LC3 reporters, p62 turnover assays, or lysosomal acid lipase inhibition), nor have we ruled out contributions from neutral lipolysis via ATGL/HSL^47^.

Although we observed upregulation of all three Rab5 isoforms (Rab5A, Rab5B, Rab5C) in HCC (Figure 6A), their individual contributions to lipid metabolism and tumor progression remain undefined. Literature suggests these isoforms differ in localization, binding partners, and possibly function^32,83^. Future studies will therefore employ isoform-specific CRISPR/Cas9 knockout or RNAi to dissect their unique roles in lipophagy and HCC growth. Equally important is identifying the molecular effectors that link Rab5 to LD-lysosome trafficking; proximity-based proteomics (APEX/BioID) on isolated LDs could map the Rab5 interactome and reveal key tethering or fusion proteins. Finally, given that our Hep3B and Huh7 cell line experiments do not fully recapitulate the tumor microenvironment, it will be essential to validate Rab5’s therapeutic potential using *in vivo* cancer HCC models.

## Acknowledgement

We acknowledge Dr. Mott Justin and my lab colleagues for their thoughtful comments during lab meetings, and we are grateful to Dr. Manesh Jain and Dr. Abhijit Aithal for access to the IncuCyte system. This work was supported by the National Cancer Institute R21 Grant R21CA279878, the NIAAA R00 Grant R00AA026877, and the NIGMS R35 MIRA Award R35GM150801 (to M.B.S.). The graphical illustrations were created with BioRender.com.

## Supporting Information Legend

1A) Western blot of constitutively active Rab5 (Q79L) and dominant negative rab5 (S34N) GTP-pulldown. To validate that the GTP-bead pulldown works perfectly.

2A) E) Western blot analysis of Rab5 and EEA1 GTP-pulldown in 48h NAP and DMSO control treated condition (nutrient rich media) in Hep3B cell. Note: decrease of Rab5 in the Rab5 inhibited condition (NAP) compared to control (Nutrient rich media).

2B) Quantification of ratio of GTP/total Rab5 in GTP-pull down assay from n=3 independent experiments. Quantification of ratio of GTP/total EEA1 in GTP-pull down assay from n=3 independent experiments. (mean ± SD, p=0.2785, **p<0.005 by unpaired two-tailed *t*-test).

