## Supplementary figures and images for "RAB5 NUCLEOTIDE BINDING PROMOTES β-OXIDATION TO FUEL HEPATOCELLULAR CARCINOMA CELL PROLIFERATION"

### Supplemental figure 1 and 2

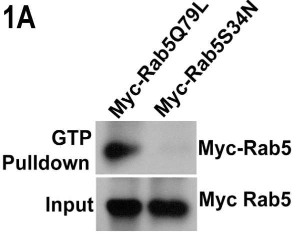

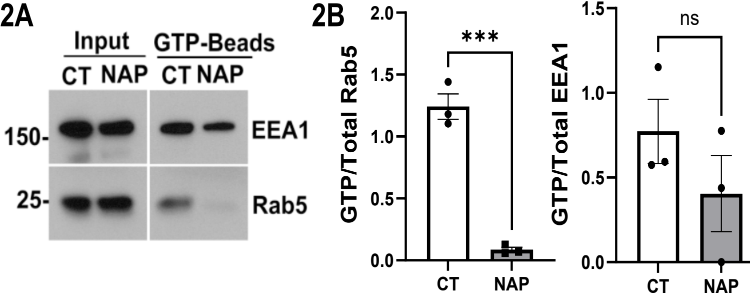
