## Supplemental Table 2 for "RAB5 NUCLEOTIDE BINDING PROMOTES β-OXIDATION TO FUEL HEPATOCELLULAR CARCINOMA CELL PROLIFERATION"

**Supplementary Table 2**

Table 2: Overrepresentation of Rab5 Isoform-Associated Genes among Cancer Hallmark Gene Sets

| **Cancer Hallmark** | **Overlap** | **P-value** | **Adjusted P-value** | **Odds Ratio** | **Hallmark vs. hallmark** |
| --- | --- | --- | --- | --- | --- |
| SUSTAINING PROLIFERATIVE SIGNALING | 3/3574 | 0.01352 | 0.02366 | 22.4 | 0.74 |
| GENOME INSTABILITY | 0/1 | nan | 1.0 | nan | 0.0 |
| EVADING GROWTH SUPPRESSORS | 3/3288 | 0.01052 | 0.02366 | 24.95 | 0.8 |
| EVADING IMMUNE DESTRUCTION | 1/749 | 0.14245 | 0.1662 | 11.42 | 1.17 |
| SUSTAINED ANGIOGENESIS | 0/1 | nan | 1.0 | nan | 0.0 |
| TISSUE INVASION AND METASTASIS | 3/2318 | 0.00369 | 0.0129 | 38.34 | 1.14 |
| TUMOR-PROMOTING INFLAMMATION | 3/769 | 0.00013 | 0.00094 | 129.97 | 3.43 |
| RESISTING CELL DEATH | 1/1941 | 0.34015 | 0.34015 | 4.04 | 0.45 |
| REPROGRAMMING ENERGY METABOLISM | 0/1 | nan | 1.0 | nan | 0.0 |
| REPLICATIVE IMMORTALITY | 1/547 | 0.10547 | 0.14765 | 15.87 | 1.61 |

This table assesses the enrichment of Rab5 isoform associated genes within the ten integrated “Hallmark” gene sets (n = 6,763) representing key cancer-related processes.
